# Latency and amplitude of the stop-signal P3 event-related potential are related to inhibitory GABA_a_ activity in primary motor cortex

**DOI:** 10.1101/2020.09.15.298711

**Authors:** Megan Hynd, Cheol Soh, Benjamin O. Rangel, Jan R. Wessel

## Abstract

By stopping actions even after their initiation, humans can adapt their ongoing behavior rapidly to changing environmental circumstances. The neural processes underlying the implementation of rapid action-stopping are still controversially discussed. In the early 1990s, a fronto-central P3 event-related potential (ERP) was identified in the human EEG response following stop-signals in the classic stop-signal task, accompanied by the proposal that this ERP reflects the “inhibitory” side of the purported horse-race underlying successful action-stopping. Later studies have lent support to this interpretation by finding that the amplitude and onset of the stop-signal P3 relate to both overt behavior and to movement-related EEG activity in ways predicted by the race model. However, such studies are limited by the ability of EEG to allow direct inferences about the presence (or absence) of true, physiologically inhibitory signaling at the neuronal level. To address this, we here present a cross-modal individual differences investigation of the relationship between the features stop-signal P3 ERP and GABAergic neurotransmission in primary motor cortex (M1, as measured by paired-pulse transcranial magnetic stimulation). Following recent work, we measured short-interval intracortical inhibition (SICI), a marker of inhibitory GABA_a_ activity in M1, in a group of 41 human participants who also performed the stop-signal task while undergoing EEG recordings. In line with the P3-inhibition hypothesis, we found that subjects with stronger inhibitory GABA activity in M1 also showed both faster onsets and larger amplitudes of the stop-signal P3. This provides direct evidence linking the properties of this ERP to a true physiological index of motor system inhibition. We discuss these findings in the context of recent theoretical developments and empirical findings regarding the neural implementation of inhibitory control during action-stopping.

## 1. Introduction

Inhibitory control of movement is a key cognitive control function implemented by the human brain, which allows humans to cancel actions even after their initiation. The neurophysiological underpinnings of inhibitory control are still subject of controversial debate. Neuroimaging studies have identified a distributed network of frontal and basal ganglia brain regions that is activated by sudden signals to stop an action (Aron & Poldrack, 2006; Chevrier, Noseworthy, & Schachar, 2007). There is now widespread agreement that the exact timing of activity within that brain network, rather than the magnitude of its activity alone, is paramount to the successful implementation of motor inhibition (W. Chen et al., 2020; Schmidt, Leventhal, Mallet, Chen, & Berke, 2013; Wessel & Aron, 2015). This notion consistent with early behavioral and theoretical work on the stop-signal paradigm, which holds that successful stopping of movement is achieved through a race between inhibitory control process(es) (triggered by a signal to stop an action) and the processes governing movement initiation and execution (Logan, Cowan, & Davis, 1984; Verbruggen & Logan, 2009). While there is no overtly observable response on successful stop trials, the assumptions of this ‘horse-race’ model allow the calculation of a latent variable – ‘stop-signal reaction time’ (SSRT) – that expresses the latency of the inhibitory process (Verbruggen et al., 2019). Shorter SSRTs reflect a faster, more efficient stop-process, which allows actions that are even further along in their initiation to be successfully stopped. With some modifications (Boucher, Palmeri, Logan, & Schall, 2007; Schmidt & Berke, 2017), this horse-race model of inhibitory control in the stop-signal task is still widely accepted today. Since a key implication of the horse-race model is that the timing of the ‘stop’-process is paramount to the success of action stopping, studying the neural underpinnings of inhibitory control necessitates the usage of methods with sufficient temporal resolution, such as electroencephalography (EEG).

Early work using EEG in healthy humans has resulted in the suggestion that the frontal-central P3 event-related potential following stop-signals may reflect the inhibitory process in the stop-signal task (Jong, Coles, Logan, & Gratton, 1990). Strong support for this notion came from subsequent work (Kok, Ramautar, Ruiter, Band, & Ridderinkhof, 2004), in which the latency of the stop-signal P3 was found to index stopping success: within subjects, the stop-signal P3 peaked earlier on successful compared to failed stop-trials. This property reflects a direct correspondence between the features of the stop-signal P3 and the predictions of the race model. However, controversy remained. Specifically, several authors argued that since the peak of the fronto-central P3 occurs after SSRT, it is unlikely that the process signified by this neurophysiological signal is critical to the success of motor inhibition (Dimoska, Johnstone, Barry, & Clarke, 2003; Huster, Enriquez-Geppert, Lavallee, Falkenstein, & Herrmann, 2013; Naito & Matsumura, 1996). A later study, however, used single-trial analyses to show that the *onset* of the fronto-central stop-signal P3 (instead of its peak) not only occurs early enough to fall within the pre-SSRT time period, but is also highly positively correlated with SSRT– i.e., subjects with a a faster onset of the stop-signal P3 also showed faster SSRT (Wessel & Aron, 2015). While this latter association has since been independently replicated (Huster, Messel, Thunberg, & Raud, 2020) the association between the stop-signal P3 and inhibitory processing still remains controversial. One argument is that additional ERPs, including ERPs that predate the P3 and likely reflect attentional processes, also correlate with SSRT (Huster et al., 2020; Skippen et al., 2020). In addition to this debate focusing on the electrophysiology of stopping, the fundamental assumptions underlying the SSRT computation have also recently been called into question themselves, with some authors arguing that SSRT systematically overestimates the latency of the stopping process – specifically because the classic SSRT computation does not take into account trials in which the inhibitory control process is not triggered at all (Matzke, Curley, Gong, & Heathcote, 2019; Patrick Skippen et al., 2019). Therefore, the notion that SSRT should be used as the ultimate benchmark to evaluate the involvement of specific neural indices in action-stopping has become more controversial itself.

In light of these recent criticisms of SSRT, some more recent neuroscientific studies have turned away from this latent behavioral variable and towards evaluating purported EEG indices of inhibitory control in direct relation to the (neuro)physiological activation of the motor system. The motor system is the final stage of the neural system in which the ‘go’-side of the horse-race underlying action-stopping will manifest, and motor activity can therefore function as a precise indicator of how much inhibitory control is or was necessary to stop a given action. Therefore, motor activity can be related to trial-to-trial estimates of potential inhibitory control signatures, and that relationship can then be scrutinized vis-à-vis the predictions of the horse-race model. Using such approaches, two recent such studies in particular have provided notable support for the classic interpretation of the fronto-central P3 as an index of inhibitory control processes. In one example, the lateralized readiness potential – an index of late-stage motor preparation observable in EEG recordings over sensorimotor cortex – was used to show that successfully stopped trials with more residual motor activity also showed larger stop-signal P3 ERPs (Wessel, 2018). In other words, successfully cancelled actions that were more advanced (and hence required more inhibitory control to cancel) were accompanied by a larger stop-signal P3. In another study, force recordings were used to provide further evidence along the same lines. Specifically, Nguyen, Albrecht, Lipp, & Marinovic (2020) found that on failed stop-trials (i.e., trials with a stop-signal in which the go-response was not successfully stopped) larger P3 amplitudes were found when the erroneous response was produced with reduced force. This suggests that trials on which the incorrectly executed response showed stronger residual signs of partial inhibition also were accompanied by larger stop-signal P3 amplitudes.

Studies like these can provide unique insights into the dynamics of action-stopping, since they do not have to rely on SSRT to evaluate the potential neurophysiological concomitants of the processes underlying the horse-race. However, even those studies cannot provide a direct proof of the proposition that the P3 (or any other EEG signature) directly relates to the physiological inhibition of the motor system. Since EEG does not offer insights into the physiological excitation or inhibition of the neuronal populations underlying the scalp signal, EEG-derived signatures such as ERPs cannot resolve whether an observed reduction of a motor process is truly due to the activity of inhibitory processes at the neural level. Therefore, to investigate whether specific EEG indices are truly related to the actual physiological inhibition of the motor system, EEG needs to be supplemented with other methods.

Notably, the combination of transcranial magnetic stimulation (TMS) and electromyography (EMG) provides a well-established method that allows this exact type of measurement. Specifically, when single pulses of TMS are applied to motor representations in primary motor cortex (M1), they result in overtly observable amplitude deflections in EMG recordings from the associated muscle. These deflections, called motor-evoked potentials (MEP), index the net cortico-spinal excitability of the associated motor tract (Rossini et al., 1994). Additionally, more advanced TMS methods make use of paired-pulse protocols to directly index the activity of local inhibitory γ-aminobutyric acid (GABA) neuron populations in M1 (Kujirai et al., 1993; Sanger, Garg, & Chen, 2001). GABA is the primary inhibitory neurotransmitter in the brain, including in M1 (Hall et al., 2011; Stagg et al., 2011). In paired-pulse TMS protocols, introducing a ‘conditioning’ pulse prior to the MEP test pulse results in a reduction of the MEP compared to an unconditioned, single-pulse MEP. Depending on the latency difference between conditioning and test pulse, this relative reduction in the MEP is either called short-interval or long-interval intracortical inhibition (SICI / LICI). SICI is typically observed when the subthreshold conditioning pulse precedes the test pulse by 1-6ms (Kujirai et al., 1993). It is thought that SICI results from the activation of the low threshold inhibitory system by the conditioning pulse, resulting in the production of inhibitory post-synaptic potentials via fast ionotropic GABA_a_ receptors (Vincenzo Di Lazzaro et al., 2006; Stagg et al., 2011). LICI, on the other hand, is observed when a suprathreshold conditioning pulse precedes the test pulse by 50-200ms (Valls-Solé, Pascual-Leone, Wassermann, & Hallett, 1992), which is consistent with the slower production of inhibitory post-synaptic potentials produced via metabotropic GABA_b_ receptors (Chu, Gunraj, & Chen, 2008; McCormick, 1989). Pharmacological studies have largely supported these ideas for both SICI (V Di Lazzaro et al., 2000; Ziemann, Lönnecker, Steinhoff, & Paulus, 1996) and LICI (McDonnell, Orekhov, & Ziemann, 2007), with some proposing distinct, but interacting, neuronal circuits as the possible origin (R. Chen, 2004; Sanger et al., 2001). SICI and LICI are reduced during volitional movements (Hammond & Vallence, 2007; Reynolds & Ashby, 1999; Ridding, Taylor, & Rothwell, 1995), and have been shown to be differentially modulated by specific motor tasks (Kouchtir-Devanne, Capaday, Cassim, Derambure, & Devanne, 2012; Opie, Ridding, & Semmler, 2015). Therefore, SICI and LICI are popular indices of the activity of local GABA_a/b_ receptors in M1 and their relationship to motor behavior.

In line with the proposition that stopping an action would involve the activity of true, physiological inhibition of the motor system, studies using the stop-signal task have also shown that SICI and LICI are related to the success of stopping. Indeed, increased SICI has been demonstrated in subjects who are required to rapidly inhibit movements (Coxon, Stinear, & Byblow, 2006), while LICI has been shown to increase when subjects possess prior knowledge of when they need to inhibit specific movements (Cowie, MacDonald, Cirillo, & Byblow, 2016; Sohn, Wiltz, & Hallett, 2002). In other words, GABA_a_ activity, indexed by SICI, appears to be increased during rapidly exerted, reactive inhibitory control following a stop-signal, while GABA_b_ neurons seem to be anticipatorily recruited during situations in which inhibitory control can be proactively deployed towards specific effectors. Indeed, in perhaps the most direct demonstration of the relationship between SICI-indexed GABA_a_ activity in M1 and stopping behavior, Chowdhury, Livesey, Blaszczynski, & Harris (2018) have recently shown that SICI measured at baseline (i.e., in the absence of a task) is directly related to SSRT across subjects. In their study, subjects with stronger SICI at baseline also showed faster SSRTs during the stop-signal task. This shows that the ability to recruit inhibitory neuronal circuitry in M1 is directly related to the ability to stop an action. While the recent criticism of SSRT as a measurement variable itself (see above) can also be levied against these findings, the results of Chowdhury and colleagues’ study do indicate that GABAergic inhibitory neural activity in M1 can potentially be used to probe an individual’s ability to physiologically inhibit the motor system – and ultimately, to stop an action. In the current study, we attempt to capitalize on this utility of SICI to directly test whether the stop-signal P3, as a purported index of inhibitory control processing in frontal cortex, is related to the physiological activity of GABA_a_ neurons in M1.

To this end, we combined the SICI approach of Chowdhury et al. (2018) with additional recordings of scalp EEG during the stop-signal task. This was done with the explicit goal of directly relating each subjects’ SICI measurement to the two core features of the fronto-central stop-signal P3 that have been proposed to index inhibitory control activity: its onset latency and its amplitude. Doing so achieves two overarching goals: First, using a direct index of the activity of inhibitory neurotransmission in M1 will allow us to evaluate whether the fronto-central stop-signal P3 ERP is related to the physiological inhibition of the motor system. Second, relating two measurements of brain activity that purportedly index inhibitory control without having to rely on a latent behavioral variable (viz., SSRT) would provide a direct validation of both measures and their purported involvement in action-stopping using overtly measurable physiological variables.

Based on the above-mentioned findings regarding the relationship between SICI, the P3, and stopping behavior, we hypothesized that subjects with stronger GABA_a_ activity would show faster onset latencies and larger amplitudes of the stop-signal P3. Furthermore, we also collected baseline estimates of LICI to test whether GABA_b_ activity is related to the behavioral implementation of proactive inhibitory control.

## 2. Materials and Methods

### 2.1 Data availability

[All data, analysis scripts, and task code will be made available on the OSF upon acceptance of the manuscript.]

### 2.2 Participants

Forty-one healthy adult subjects participated in this study (27 female, 14 males; mean age: 21.6 years, SD: 3.85, range [18 30] ; all right-handed). Subjects were recruited either through UI’s Department of Psychological and Brain Sciences sign-up system or through a mass recruitment email. Subjects recruited were granted either class credit for their participation or were compensated with $15/hr. All procedures were approved by the University of Iowa’s Institutional Review Board (#201612707, #201511709). Prior to experimentation, all subjects were screened for abnormal (non-corrected) vision or hearing, as well as for contraindications of TMS (Rossi, Hallett, Rossini, Pascual-Leone, & Group, 2009).

### 2.3 Stimulus presentation

All stimuli were presented on a 19-inch Dell flat screen monitor connected to an IBM-compatible PC running Fedora Linux and MATLAB 2015b. Stimuli were presented using Psychtoolbox 3 (Brainard, 1997). Responses were made using a standard QWERTY USB keyboard.

### 2.4 Experimental task

Our variant of the stop-signal task alternated between blocks of the classic stop-signal paradigm and blocks of a pure-go task (which was identical in layout and timing to the stop-signal blocks but did not include any stop-signals, ***Figure 1***). Pure-go blocks were included in this study to measure proactive control behavior (i.e., reaction time slowing on go-trials in stop-vs. pure-go blocks), in addition to reactive control (Chikazoe et al., 2009; Jahfari, Stinear, Claffey, Verbruggen, & Aron, 2010; Verbruggen & Logan, 2009). This was done to explore the proposed association between LICI and behavioral indices of proactive inhibitory control (Cowie et al., 2016), a secondary goal of the current study.

**Figure 1.**
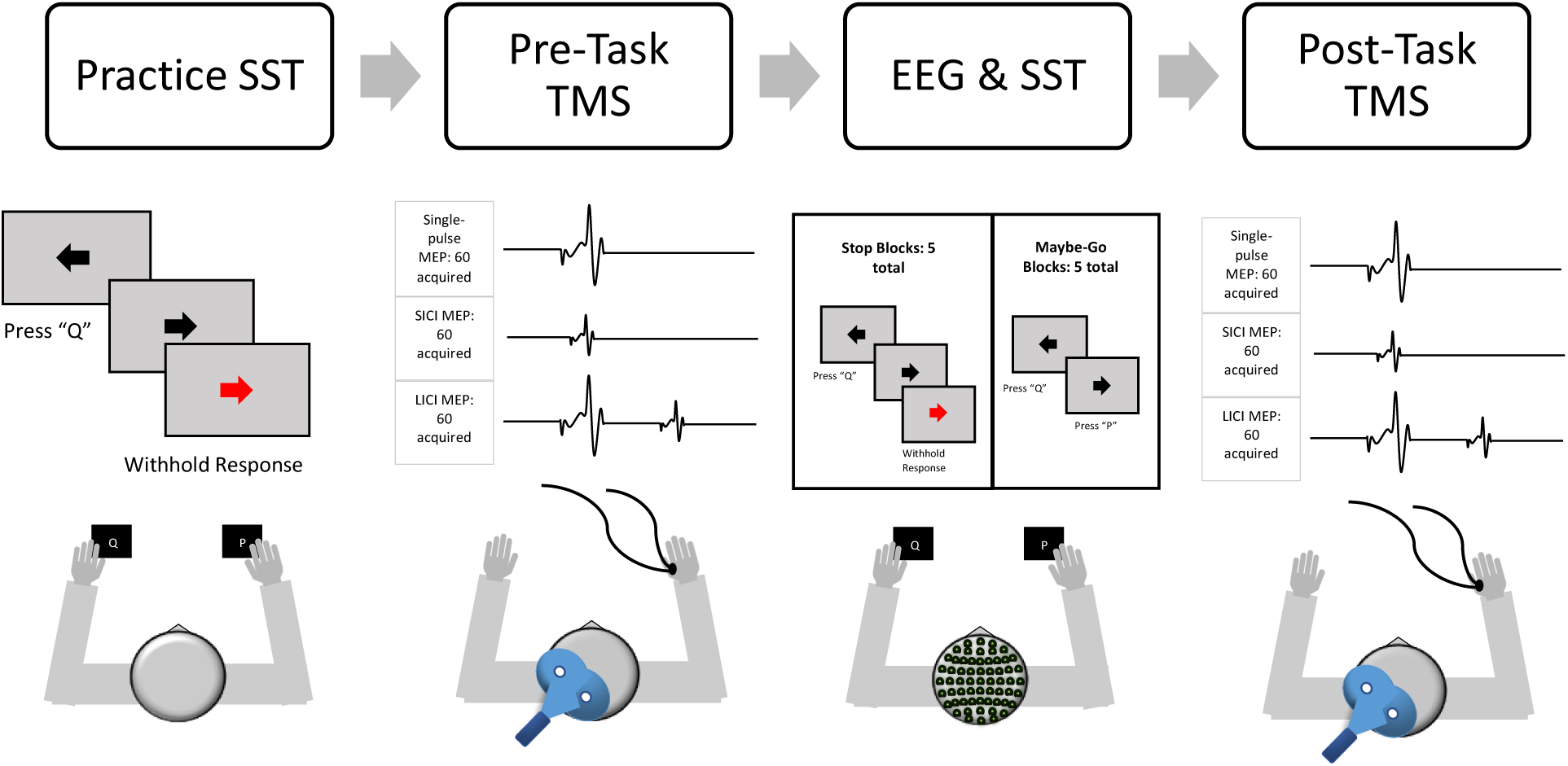
Study design, task diagram, and illustration of the TMS and EEG procedures.

Stimuli were presented on a gray background. Each trial began with a black fixation cross (500ms), followed by a black arrow (go-signal) pointing either left or right, displayed for 1,000ms. Subjects were instructed to press the “q”-key on the keyboard with their left index finger in case of a left facing arrow and “p” with their right index finger in case of a right facing arrow. Responses were to be made within the 1,000ms window during which the stimulus was presented on the screen. Each trial was followed by a 1,500ms inter-trial interval (ITI). If no response was made during the response window, the first 1,000ms of the ITI included a red “Too Slow!” message presented on the screen. In stop-signal blocks, a stop-signal (i.e., the black go-signal arrow changing to red color) was presented after the go-signal on 25% of trials. The stop-signal delay was initially set to 200ms and was then subsequently adjusted in steps of 50ms (which were added to the stop-signal delay after successful stop-trials and subtracted after failed stop-trials) with the goal of achieving an overall p(stop) of approximately 0.5. Participants completed 2 blocks of practice with the stop-signal task, and then performed 10 total blocks, alternating between stop-signal blocks (“There will be stop-signals. Responding quickly on go-trials and stopping successful on stop-trials are equally important.”) and pure-go blocks (“There will be no stop-signals. Respond as fast as possible.”). We altered the type of the first block after each subject to counterbalance the order. Two blocks were removed from one participant who was pressing the wrong buttons towards the end of the experiment. To achieve a balanced number of go-trials from each task context, each block contained 48 go-trials. In addition, the stop-signal blocks contained 16 stop-trials. In total, this resulted in 240 go-trials from pure-go blocks, 240 go-trials from stop-signal blocks, and 80 stop trials per subject.

### 2.5 Procedure

Each subject participated in a one-time experimental visit, lasting approximately 3 hours, which consisted of a) practicing the task, b) pre-task measurements of SICI/LICI, c) EEG recording during the task, and d) post-task measurements of SICI/LICI (***Figure 1***).

### 2.6 TMS Protocol

Each subject had their right hand cleaned using alcohol wipes and an abrasive pad, before two Ag/AgCl electrodes were adhered parallel to the belly of the first dorsal interosseus (FDI) muscle, while a third electrode (reference) was placed over the ulnar head. Electrodes were connected to a Grass P511 amplifier (1000 Hz sampling rate, filters: 30 Hz high-pass, 1000 Hz low-pass, 60 Hz notch). Amplified data were sampled via a Micro 1401-3 sampler (Cambridge Electronic Design) and visualized/recorded using Signal software (Version 6; Cambridge Electronic Design).

Each subject adorned a Lycra cap prior to TMS stimulation to mark TMS coil locations during hotspotting. TMS stimulation was performed using a MagStim BiStim^2^ system with a 70mm figure-of-eight coil. The coil was held 45 degrees to the coronal plane, over the left primary motor cortex (M1) of each subject, inducing an anterior-posterior (AP) current (see Figure 2). Hotspotting was performed while maintaining coil angle, in order to locate the locus of optimal FDI stimulation, which was then marked on the Lycra cap. Resting motor threshold (RMT) was then defined as the minimum intensity required to induce MEPs of amplitudes exceeding 0.1 mV peak to peak in 5 of 10 consecutive probes (Rossini et al., 1994).

**Figure 2.**
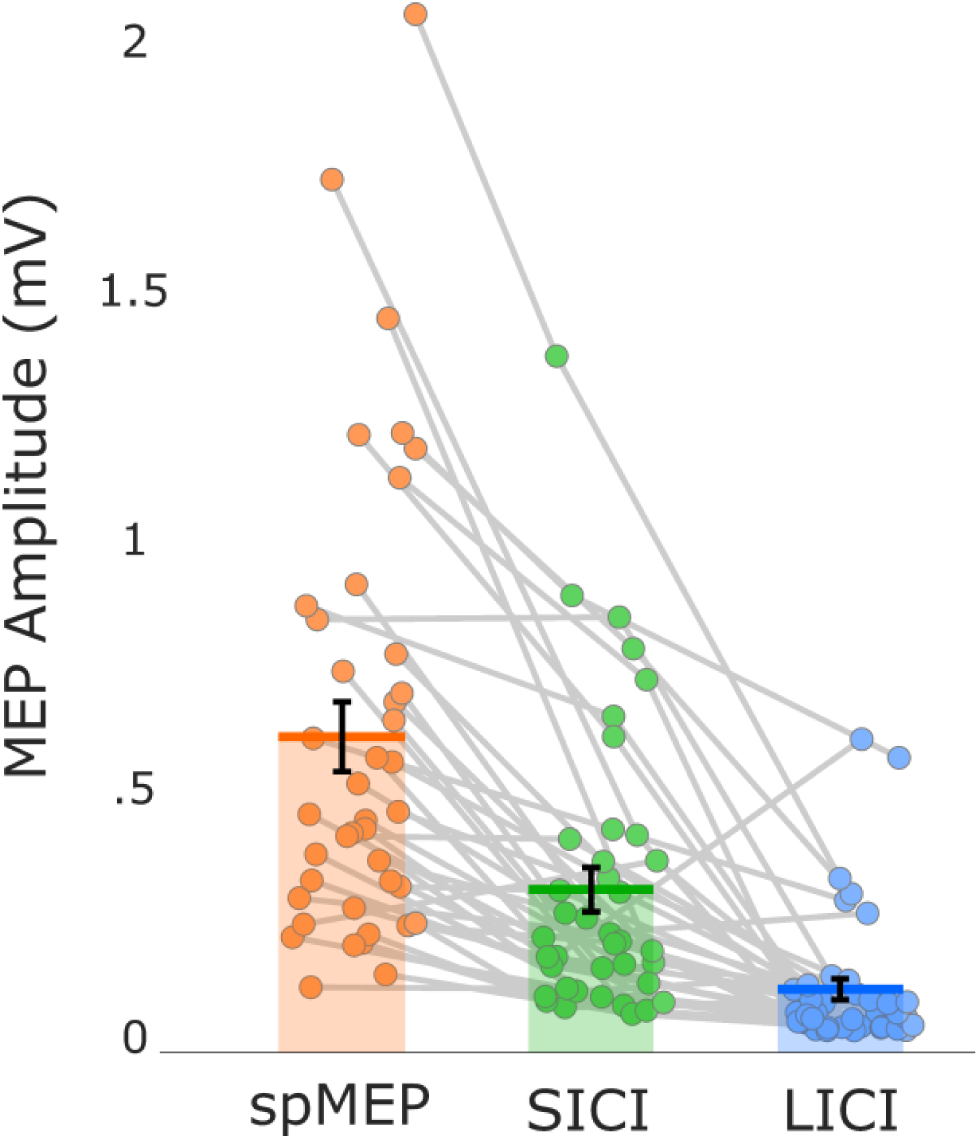
Average motor evoked potentials from all three stimulation conditions. Orange: single pulse unconditioned MEP (spMEP). Green: short-interval intracortical inhibition (SICI). Blue: long-interval intracortical inhibition (LICI). Horizontal lines represent the group mean, the confidence interval represents the standard error of the mean.

### 2.7 Short-/ Long-Interval Intracortical Inhibition protocols

All TMS/EMG data were collected while the participant was at rest. We aimed to quantify both LICI and SICI. Hence, data were collected in 9 blocks of 20 trials; with 3 blocks dedicated to each measurement (i.e., 3 blocks each of paired-pulse SICI, paired-pulse LICI, and the single-pulse, unconditioned MEP). These blocks were acquired in randomized order. For SICI, the conditioning pulse was sent at 80% RMT and the test pulse sent at 120% RMT, with the inter-stimulus interval set at 2ms (Peurala, Müller-Dahlhaus, Arai, & Ziemann, 2008). For LICI, both the conditioning and test pulse were sent at 120% RMT with 100ms ISI (Sanger et al., 2001). Unconditioned, single-pulse MEPs were elicited with a single-pulse at 120% RMT. All MEPs were screened after collection using in-house MATLAB software. Trials were excluded if the root mean square of the pre-stimulus baseline EMG exceeded 0.1mv or if the peak amplitude of the MEP was outside the range of -2.99mV -2.99mV. SICI and LICI were computed by using a following equation:

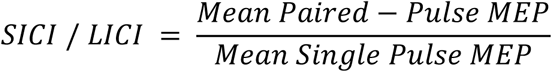

When comparing the amplitudes of the MEP before and after the task, it became evident that there was a significant difference between the pre- and post-task MEP amplitudes (post-stim MEPs larger; t(40) = 2.86, p < .01). Therefore, we decided to focus our investigation on the pre-task measurements. This was done for two reasons: First, alertness and sleep pressure may affect motor excitability (Gennaro et al., 2007; Noreika et al., 2019). Given that the post-task TMS measurements were collected after an extended period of performing a repetitive behavioral task, those measurements were likely to be affected by these confounding variables, potentially explaining the discrepancy. Second, inhibitory control training itself can also change the amplitude of the MEP in the involved muscles (Majid, Lewis, & Aron, 2015). Therefore, we believe the pre-task measurements to be a more accurate representation of GABA activity, unaffected by both fatigue and potential inhibitory control training effects.

### 2.8 Behavior Analysis

Within the stop-blocks, go- and failed stop-trial reaction time were compared to ensure that the requirements of the horse race-model were met (specifically, failed stop reaction time had to be faster than go-trial reaction time, (Logan et al., 1984). Furthermore, p(stop) was investigated to ensure that the stop-signal delay staircase algorithm was effective in achieving an approximate stopping success rate of .5. SSRT was calculated from the stop-block data using the integration method with replacement of go-trial omission errors (Verbruggen et al., 2019). Lastly, to explore the purported relationship between LICI and proactive control behavior, we also computed the response delay effect between stop-signal and pure-go blocks using the following equation:

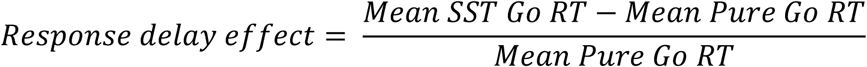

### 2.9 EEG recording

EEG data was recorded at a sampling rate of 500Hz (10s time-constant high-pass and 1000 Hz low-pass hardware filter) using a 64-channel active electrode cap connected to an actiCHamp amplifier (BrainProducts, Garching, Germany). The reference electrode was Pz and a ground electrode was placed at Fz.

### 2.10 EEG preprocessing

EEG data were preprocessed using MATLAB functions and the EEGLab toolbox. Raw EEG data were bandpass filtered (.5 – 50 Hz) using a Hamming windowed sinc FIR filter. Following filtering, all participants’ continuous EEG data were visually inspected and any segments that included non-stereotypical artifacts were removed. Data were then re-referenced to the common average and submitted to a temporal infomax ICA decomposition algorithm (Bell & Sejnowski, 1995). The resulting component matrix was visually inspected in order to identify independent components (ICs) that reflected stereotypical artifacts (blinks and saccades), which were removed from the data by means of selective backprojection.

### 2.11 P3 onset calculation

The procedures to calculate the onset of the P3 ERP in each participant were designed to match the methodology of Wessel & Aron (2015). First, one IC was identified from each participant’s ICA solution to represent the fronto-central P3. In order to select this IC, we manually identified candidate ICs that showed maximal IC weights around the fronto-central electrodes (Cz, FCz) in the scalp topographies. We then back projected these candidate ICs into the channel space to compute ERP using epochs time-locked to stop-signal onsets in order to confirm that there was N2/P3 following stop-signals. Using this procedure, we identified a single IC for each participant that fulfilled both criteria.

Subsequently, to validate that these chosen ICs showed the previously demonstrated properties of the P3 in relation to behavior in the SST, we tested a) if the onset of the P3 was faster in successful compared to failed stop trials within subjects and b) whether the successful stop P3 onset correlated with SSRT across subjects. To this end, the thusly selected IC was back projected into the channel space and its properties were investigated in the SST blocks. Four types of events were time-locked (−300 – 700ms) for each epoch: successful stop (SS), failed stop (FS), and go trials paired with each type of stopping. Each stop trial was paired with a go trial in which a pseudo-event was generated at the same SSD. This approach allowed us to compare the stop-related activity with matching time ranges on go trials (Wessel & Aron, 2015). Within each subject, pairs of stop and matched go trials were compared for each sample point using label shuffling permutation testing (10,000 iterations, p = .01, corrected for multiple comparisons using the false-discovery rate method, FDR, (Benjamini, Krieger, & Yekutieli, 2006). The onset of P3 was identified by locating the P3 peak latency first, then moving ‘backwards’ (i.e., towards the stop-signal) until there was no more significant difference between stop and matching go trials. We then compared the P3 onsets of SS vs. FS trials using a paired-samples t-test. The correlation between the SS P3 onsets and SSRT estimates was tested using Pearson’s product-moment correlation coefficient.

### 2.12 Brain-behavior and brain-brain correlations

All correlations were tested using Pearson’s correlation coefficient. To test our main hypothesis, SICI was correlated with the onset latency of the stop-signal P3 across subjects. We predicted a positive correlation, signifying that participants with stronger GABA_a_ activity (i.e., lower SICI values) would also show faster neural stopping activity (i.e., lower P3 onset values in msec after the stop-signal).

Moreover, we correlated each subject’s SICI measurement with the amplitude of the trial-averaged ERP of the selected IC, which was quantified for each sample point in the 700ms time period immediately following the stop-signal. This was done to investigate a potential relationship between the amplitude of the P3 (and potential additional fronto-central ERPs represented in the same IC), without making prior assumptions about the latency of the P3 peak. These analyses were performed separately for both the successful and failed stop-trial waveforms.

Furthermore, we attempted to investigate two previously reported associations between SICI/LICI. First, aimed to replicate the previously reported correlation between baseline SICI and SSRT (Chowdhury et al., 2018). Second, we correlated the response delay effect (i.e., the purported behavioral marker of proactive inhibitory control) with the LICI measurement, as previous research has shown that LICI strength is modulated by task related expectations of stopping (Cowie et al., 2016).

## 3. Results

### 3.1 Behavior

In the stop-signal blocks, all participants showed slower Go RT (mean: 554 ms; SD: 119) compared to failed stop trial RT (mean: 488 ms; SD: 114), indicating all data were consistent with that assumption of the horse race model. Mean p(stop) was .53 (SD: .05; .45 – .71) which validated the effectiveness of the staircase procedure. Mean SSRT was 237 ms (SD: 31). The mean response delay effect between stop-signal and pure-go blocks was .41 (SD: .27; mean pure go RT: 393 ms, SD: 50; mean SST go RT: 554 ms; SD: 119).

### 3.2 SICI & LICI

Across all subjects, the mean unconditioned single-pulse MEP amplitude was .60 (SD: .45). The SICI conditioned paired-pulse MEP was .30 (SD: .29), which presents a significant reduction from the single-pulse MEP (t(40) = 6.46, p = 5.33 * 10^−8^), reflecting the presence of SICI. The mean reduction of the MEP due to SICI conditioning pulses was 48.6 %, and SICI was numerically present in 39 out of 41 subjects (nominal SICI MEP/MEP ratio < 1). The average LICI conditioned paired-pulse MEP was .10 (SD: .13), which presents a significant reduction from the single-pulse MEP (t(40) = 7.68, p = 1.07*10^−9^), reflecting the presence of LICI. The mean reduction of the MEP due to LICI conditioning pulses was 82%, and LICI was found in 40 out of 41 subjects (nominal LICI MEP/MEP ratio < 1). The MEP results can be found in Figure 2.

### 3.3 Validation of P3 component selection

Replicating Wessel & Aron (2015), the P3 onset on successful stop-trials occurred significantly earlier compared to failed stop-trials (t(40) = 7.17; p < .001, Cohen’s d = 1.12) and was positively correlated with SSRT (r = 0.35, p < .05; one participant removed from sample due to Cook’s distance > 4/N). These properties, alongside the grand-average ERP waveform of the selected components, can be found in ***Figure 3***.

**Figure 3.**
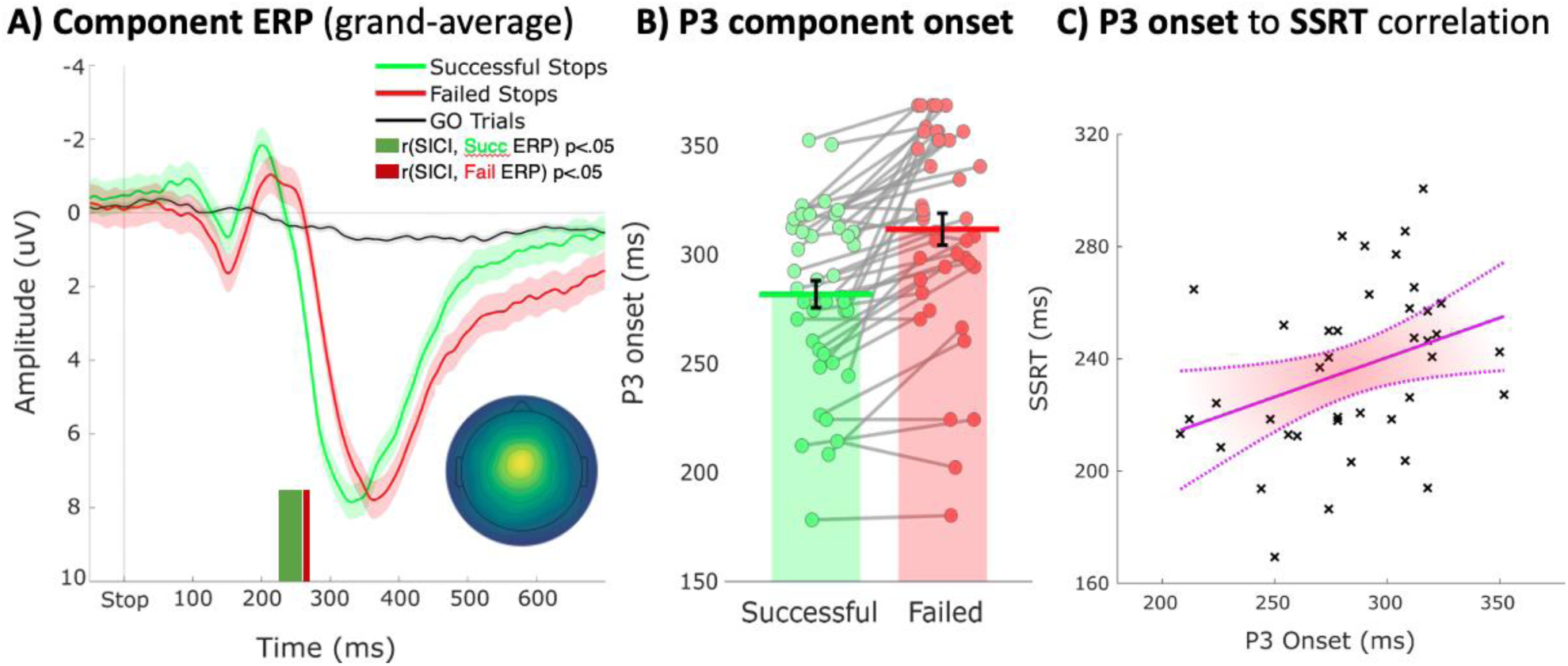
Validation of the selected component reflecting the stop-signal P3 ERP. A) Grand-average of the back-projected channel ERP at fronto-central electrodes (Cz and FCz), separately for successful and failed stop-trials. Inset: Mean component topography (rectified at Cz /FCz). Green and red bars on the bottom of the plot highlight significant periods in which the ERP amplitude correlated with SICI. B) Results of the single-trial based P3 onset identification. P3 onset was significantly earlier for successful vs. failed stop-trials. C) Positive correlation between P3 onset latency (in ms) and SSRT (in ms), alongside confidence interval.

### 3.4 SICI – P3 correlations

In line with our primary hypothesis, SICI and the onset of the fronto-central stop-signal P3 were positively correlated (r = .37, p = .017) – in other words, subjects that showed greater GABA_a_ activity in M1 (lower SICI values) also showed faster fronto-central P3 ERPs (lower P3 onset values, ***Figure 4***). Moreover, our sample-to-sample amplitude analysis showed that SICI was correlated with stop-related activity of the selected independent component on both successful (significant time period: 228 – 260 ms after the stop-signal) and failed stop-trials (significant time period: 264 – 268 ms after the stop-signal; ***Figure 3A***). No other time periods showed a significant correlation with SICI (neither in the selected independent component, nor in the ‘regular’ ERP that was reconstructed using all non-artifact independent components).

**Figure 4.**
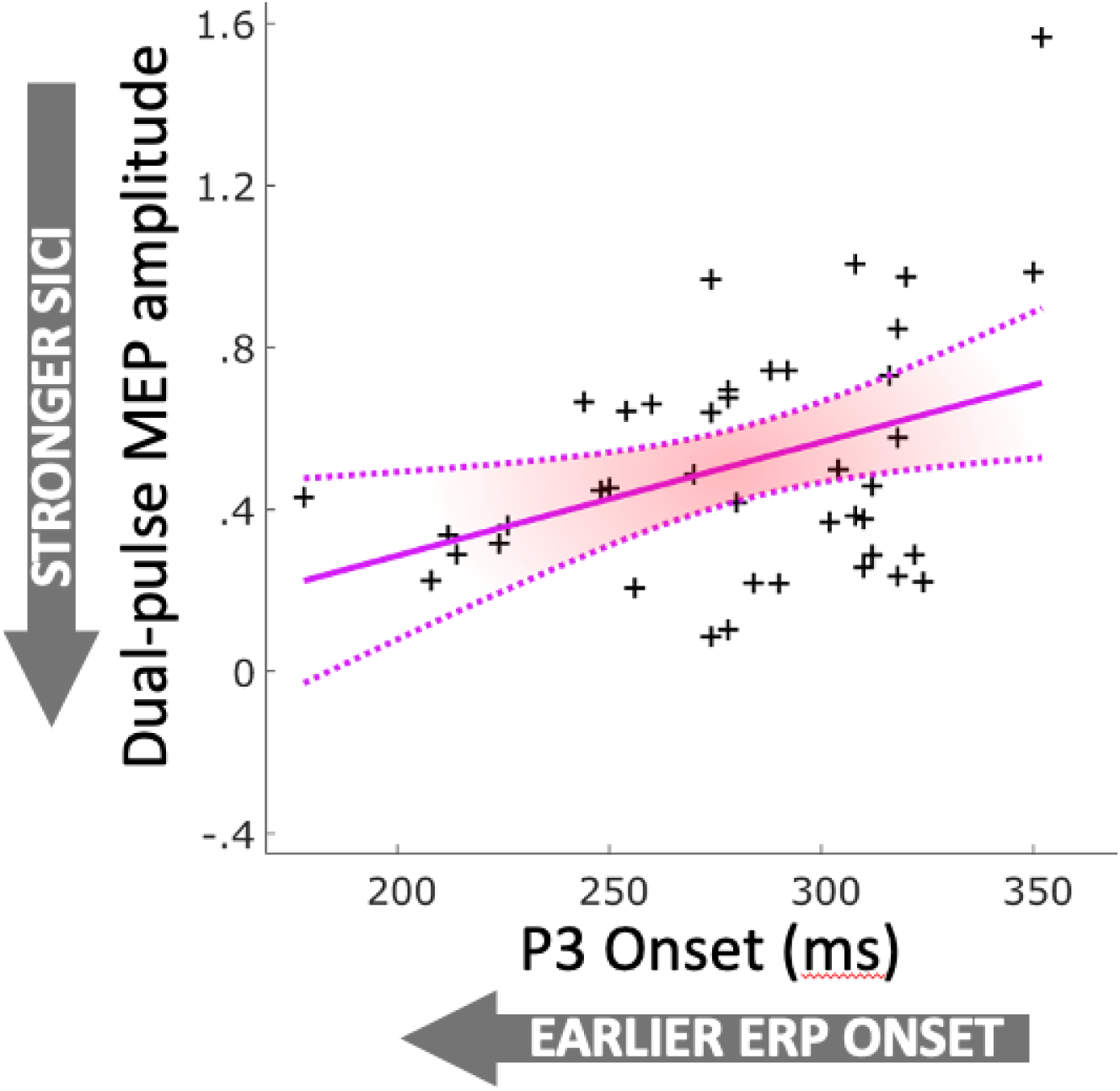
P3 onset and SICI are positively correlated across subjects; subjects with faster P3 onset (lower ms values) also show stronger SICI effects (lower dual-pulse MEP amplitudes). Magenta solid line shows best fit, dotted line shows the confidence interval.

### 3.5 SICI / LICI – behavior correlations

Contrary to our expectations, we did not find any relationship between SICI / LICI and behavior. Specifically, we did not find a significant correlation between SICI and SSRT (r = .02, p = .92). Moreover, we did not find a significant relationship between LICI and the proactive response delay effect (r = .21, p = .19).

## 4. Discussion

In the current study, we combined EEG recordings and paired-pulse TMS to investigate the relationship between a physiological indicator of GABA_a_ activity in primary motor cortex (SICI) and a purported neural index of an inhibitory control process underlying action-stopping (the fronto-central stop-signal P3). Based on previous findings that suggest a relationship between a subject’s baseline M1 GABA_a_ activity and their ability to stop an action (Chowdhury et al., 2018; Coxon et al., 2006; Sohn et al., 2002), as well as on the purported role of the fronto-central P3 as a neural index of cortical processes underlying this ability (Jong et al., 1990), we hypothesized to find a direct cross-subject correspondence between SICI and the properties of the P3 to stop-signals. This hypothesis was confirmed: participants with stronger GABA_a_ activity in M1 also showed faster and stronger activations of the fronto-central P3.

These results indicate that there is a direct relationship between the inhibitory activity of GABA_a_ neurons in primary motor cortex and the cortical processes that purportedly index inhibitory control during action-stopping. Notably, correlation does not imply causation, and indeed, our interpretation of this finding does not purport the existence of a direct causal relationship between both signals. Instead, we surmise that both stronger local GABA activity in M1 (reflected by SICI), as well as more efficient inhibitory control-related fronto-central cortical processing after stop-signals (reflected in a faster and larger P3 ERP) indicate a superior entrainment of an interconnected network of regions underlying inhibitory control of action, which includes cortical regions, inhibitory neurons in the motor system, and the subcortical basal ganglia (Aron & Poldrack, 2006; Aron, Robbins, & Poldrack, 2014; Verbruggen & Logan, 2008; Wiecki & Frank, 2013). We believe that our results suggest that subjects who generally show a faster / stronger response of the cortical systems purportedly underlying the triggering of inhibitory control processes after signals to stop an action also show stronger inhibitory signaling in motor cortex. This would indicate a superior functioning of the distributed inhibitory motor control network in these subjects, either as a result of practice, due to genetic factors, or as a result of both. Indeed, previous research has shown that the ability to exert motor inhibition can both be improved through repeated practice (Manuel, Bernasconi, & Spierer, 2013) and is partially attributable to genetic variables (Colzato, Wildenberg, Does, & Hommel, 2010; Weafer et al., 2017). Since it is broadly agreed upon that the precise timing of inhibitory control is paramount to action-stopping – especially in tasks like the stop-signal task, which purportedly reflect a race between response preparation and inhibitory control – it is likely that the individual components of the neural network underlying inhibitory control (cortical, subcortical, and motor system level) work in tight accord with one another, resulting in subjects with superior stopping abilities showing indicators of better neural functioning on several levels of measurement. Hence, humans with superior abilities to implement inhibitory control likely do so through a concerted effort that results in a tight relationship between cortical, subcortical, and motor system signatures of inhibition. However, the magnitude of the observed correlation also suggests that there is ample amount of variance that is not accounted for, suggesting that there is also variability in the functioning of the individual regions and mechanisms that work in concert to effect action-stopping. Indeed, despite the correlation observed here, some subjects may rely more on stronger inhibitory GABA networks in M1, whereas others may rely more on the frontal cortical mechanisms that ostensibly trigger the cascade of processing that ultimately results in successful action-stopping (as well as subcortical aspects that are not captured here).

Crucially, the current findings directly link the fronto-central P3 to a physiological marker of motor system inhibition in M1. Recent studies have suggested that the fronto-central P3 is related to the inhibitory demand of individual trials in which actions were successfully stopped, indicated by the activity found in (pre)motor cortex (Wessel, 2018), as well as to reductions of the force with which responses were made on unsuccessful trials (Nguyen et al., 2020). While these studies suggested a direct correspondence between motor system activity during stopping and the fronto-central stop-signal P3, they had to rely on an indirect inference to conclude that the processes reflected in the fronto-central P3 relate to the ‘true’ inhibition of motor activity. The current investigation goes beyond these studies, but corroborates their interpretation: here, the process(es) reflected in the fronto-central P3 were shown to be directly related to the physiological inhibition of the motor system.

However, a key question regarding the role of the process reflected in the P3 remains unanswered: more and more studies are showing that functional signatures of motor inhibition (including at the level of the motor system and primary motor cortex itself) occur with much faster latency after stop-signals than would be indicated by, for example, SSRT. Indeed, EMG and MEP recordings show that the first signs of motor-system inhibition in these measures emerge as early as ∼150ms after the onset of a stop-signal (Jana, Hannah, Muralidharan, & Aron, 2020; Raud & Huster, 2017). In line with this, recent studies have shown that cortical signals that are related to stopping success can also be found in similar time ranges (Huster et al., 2020; Jana et al., 2020; Skippen et al., 2020; Wessel, 2019). Just like the beforementioned EMG and MEP measurements of functional motor inhibition, these cortical signatures often precede both SSRT, as well as the typical onset latency of the fronto-central P3, by several dozens of milliseconds.

This begs the question of the role of the P3, which likely emerges too late to be causally underlying the implementation of the reported reductions of MEP and EMG at the 150ms mark. This is aspect is particularly important in the context of the current set of results. This is because the GABAergic neurons whose activity is captured by SICI are purportedly underlying the early signatures of motor system inhibition in the MEP and EMG. Given the recency of these findings, theoretical developments and empirical studies are still ongoing in this domain. However, it is notable that low-latency reductions of motor cortex such as the above described (including MEP suppression at 150ms following the stimulus) are found not just after stop-signals, but indeed after any sort of surprising (Dutra, Waller, & Wessel, 2018; G Novembre et al., 2018; Giacomo Novembre et al., 2019; Wessel & Aron, 2013) or even merely infrequent events (Iacullo, Diesburg, & Wessel, 2020). This could suggest that instead of a single, monolithic mechanism, stopping is implemented by a multi-step sequence of inhibitory processes, similar to what was proposed in the “pause-then-cancel” model by Schmidt, Berke and colleagues (Schmidt & Berke, 2017; Schmidt et al., 2013). According to this model, action-stopping includes two processes: an initial, rapid, broad ‘pause’ process, which temporarily halts all ongoing actions with the lowest latency possible, followed by a more measured, selective ‘cancel’ process, which entirely shuts off the currently invigorated motor program. Along those lines, we tentatively propose that the initial ‘pause’ process occurs after any meaningful infrequent, salient, or surprising event, resulting in the low-latency reductions of the MEP (and EMG) that have recently been reported both inside and outside of the stop-signal paradigm. During outright action-stopping in the stop-signal task, the ‘pause’ is then followed (or accompanied) by the slower ‘cancel’ process, which is more specific to stimuli that explicitly instruct the outright cancellation of ongoing action (such as a stop-signals). While we are still fleshing out the according theoretical framework in humans, we here tentatively propose that the fronto-central P3 – which purportedly originates from medial frontal regions involved in motor planning, such as the pre-SMA (Enriquez-Geppert, Konrad, Pantev, & Huster, 2010) – may reflect the second, ‘cancel’ stage (which could more broadly be understood as a reconfiguration of the active motor program). While the current study was not designed to test this interpretation, the results do suggest that both the inhibition of movement at the level of the motor system and the frontal cortical activity reflected in the fronto-central P3 are closely related – in line with the proposal that both underlying processes contribute to successful action-stopping.

While the SICI-EEG relationship in the current study lead to the confirmation of the primary hypothesis, our results notably did not confirm a secondary hypothesis and failed to replicate some prior work. Specifically regarding the latter, Chowdhury et al. (2018) had reported a positive relationship between SSRT and SICI measured at baseline – representing the main finding that motivated the hypothesis of the current study. While we did find a positive correlation between SSRT and P3 onset (replicating previous work on the EEG-behavior relationship; cf. Huster et al., 2020; Wessel & Aron, 2015), and while P3 onset was in turn correlated with SSRT as well (the main result of the current study), SICI and SSRT themselves were *not* correlated in our dataset. There are several potential reasons for this. First, as mentioned in the Introduction, SSRT has several weaknesses as a variable. In addition to the conceptual criticisms that have recently been levied against SSRT, there is also substantial variability in the algorithmic implementation of SSRT calculations. Indeed, while both Chowdhury et al. and the current study used the integration method (Verbruggen, Chambers, & Logan, 2012) to calculate SSRT, our study implemented a newly suggested procedure in which missing go-trial reaction times due to absent responses (or responses made after the response deadline) were replaced by the slowest validly counted reaction time for that same subject. This “go-omission replacement” technique has been suggested in a recent paper that was not available at the time of the Chowdhury et al. study (Verbruggen et al., 2019). Second, there were two notable ways in which our SICI measurement deviated from Chowdhury and colleagues’ work as well. First, our SICI estimates were collected *before* the performance of the task, whereas Chowdhury and colleagues measured SICI only *after* subjects performed the stop-signal task. In the current dataset, we found that MEPs collected before and after the task differed significantly, which is in line with the purported effects that both tiredness (Gennaro et al., 2007; Noreika et al., 2019) and inhibitory control training (Majid et al., 2015) have on motor excitability. We chose to omit the post-task TMS measurements for this reason, as the pre-task measurements should be unaffected by these variables (though we note that an exploratory post-hoc analysis of the post-task TMS measurements – i.e., a closer replication of Chowdhury et al.’s procedure – also revealed no significant correlation to SSRT; r = -.04, p = .78). Second, our SICI measurement procedure itself differed from the exact implementation of Chowdhury and colleagues. Specifically, while Chowdhury et al. (2018) used an inter-stimulus interval of 3ms between the conditioning pulse and the test pulse, we here used an interval of 2ms. This interval was chosen because it has been shown to reliably evoke SICI (Kujirai et al., 1993), while temporally avoiding unwanted I-wave facilitation (R. Chen, 2004; R. Chen & Garg, 2000; Hanajima et al., 2002; Peurala et al., 2008). It is entirely possible that the two procedures capture subtly different parts of the underlying GABA_a_ activity (Fisher, Nakamura, Bestmann, Rothwell, & Bostock, 2002; Peurala et al., 2008) leading to the difference in findings regarding the SICI-SSRT correlation. Finally, our implementation of the stop-signal task itself also differed from Chowdhury et al. While Chowdhury and colleagues used the classic SST implementation as programmed in the STOP-IT toolbox (Verbruggen, Logan, & Stevens, 2008), we collected SST blocks in alternation with pure-go blocks, in which no stop-signals were presented. This was done to test a secondary hypothesis about the potential role of LICI in the implementation of proactive inhibitory control (see Results). This may have affected behavior in the SST blocks in a way that influenced SSRT. Either way, our study was designed to explicitly exclude SSRT as a meaningfully variable from our comparisons, given that we aimed to directly use signatures of physiological inhibition of the motor system to evaluate the relationship of the fronto-central P3 to motor inhibition. Therefore, our study was not optimized to directly replicate the purported SICI-SSRT association demonstrated by Chowdhury et al. (2018) and we believe that our non-replication does not bear on the validity of those results (which the authors have since conceptually replicated using an event-related SICI protocol, cf. Chowdhury, Livesey, & Harris, 2019).

In regards to the current study’s secondary hypothesis, we also did not find a positive association between LICI at baseline and the degree to which participants slowed down their responding when anticipating potential stop-signals (in the SST blocks) compared to when they respond without restraint (in the pure go blocks). This analysis was originally included for reasons of convenience (i.e., because of the ease of implementation of this secondary hypothesis into the primary investigation). However, it is notable that the measurement of LICI at baseline and its comparison to the response delay effect is a substantial deviation from the original study that had purported that GABA_b_ neurotransmission in M1 may be related to the implementation of proactive control. Indeed, the proposition comes from a study by Cowie and colleagues, in which they measured LICI during a bimanual response inhibition task (Coxon, Stinear, & Byblow, 2007), demonstrating larger inhibitory effects during blocks with stop-trials (compared to only go-trial blocks) and which were positively correlated with the response delay of the left hand, following response cancellation in the right hand (Cowie et al., 2016). This suggests that LICI could reflect the tone of inhibitory circuits in M1, which is increased during tasks that may require rapid action cancellation; although this is not always the case (Cirillo, Cowie, MacDonald, & Byblow, 2018). While the current study shows that LICI at baseline does not relate to the specific behavioral phenomenon of the response delay effect, other studies (especially those that measure LICI during event-related task periods in which proactive control is exerted, rather than at baseline) may well find different results.

## 5. Conclusion

In summary, we here used a unique and novel combination of paired-pulse TMS and scalp EEG to directly investigate the relationship between inhibitory neuronal activity in primary motor cortex and control signals purportedly underlying the inhibitory control of action in frontal cortex. In line with our predictions, we found that local GABA_a_ activity in primary motor cortex was directly related to both the latency and amplitude of the fronto-central P3 ERP. This suggests that there is a direct correspondence between the properties of this frontal control signal, which has been purported to reflect a process related to the ‘horse-race’ underlying action-stopping in the stop-signal task, and the ability of subjects to physiologically inhibit motor signals. These findings provide further evidence towards the relationship between the fronto-central P3 and inhibitory control – though recent questions about the relative timing of processes remain, and will have to be addressed through sustained theoretical and empirical work.

## Acknowledgments

The authors would like to thank Nathan Chalkley and Brynne Dochterman for help with data collection. This work was supported by the National Institutes of Health (NIH R01 NS117753), the Iowa Center for Research by Undergraduates, and the Iowa Neuroscience Institute Summer Scholars’ program.

## Notes

### Competing Interest Statement

The authors have declared no competing interest.

